# ^1^H-MRS characterization of metabolic changes induced by photoperiod in the sheep brain: a promising eye into in-vivo adult neurogenesis?

**DOI:** 10.1101/2022.10.09.511460

**Authors:** Nathalie Just, Pierre Marie Chevillard, Martine Batailler, Jean-Philippe Dubois, Pascal Vaudin, Delphine Pillon, Martine Migaud

## Abstract

The hypothalamus is a central structure of the mammalian brain, which controls physiological, endocrine and metabolic brain homeostasis. Despite this essential role, little is known about the normal and altered neurochemical changes occurring within the hypothalamic structures. Here, the metabolism of the hypothalamus of ewes was investigated at 3T using proton magnetic resonance spectroscopy (^1^H-MRS). We used the sensitivity of these animals to photoperiod i.e. the ratio of day to night length, to investigate the hypothalamic metabolic changes and their relationship to hypothalamic adult neurogenesis. A longitudinal study involving 4 ewes per timepoint was conducted at 4 time points (P01, P02, P03 and P04) during long days (LD) and 4 time points during short days (SD). Significant metabolic changes were found between LD and SD at all time points in particular for glutamate (Glu), glutamine, myo-inositol and total N-acetyl-Aspartate (NAA). During SD, glutamate and glutamine concentrations were significantly smaller at P01 compared to all other time points while significant neurochemical changes occurred during the entire LD period. Neurochemical changes relative to P01 remained stable during LD and SD except for Glu and Gln which increased between P01 and P02 during SD. Relative metabolic changes were significantly higher on average for NAA and Glu and significantly smaller on average for total choline during SD compared to LD, respectively, paralleling the average changes in the numbers of neural stem cells and glial and oligodendrocyte progenitors found by immunohistochemistry. Despite important differences between MRS and immunohistochemistry in terms of spatial resolution, both techniques suggest complementary findings that should contribute to a better characterization of the hypothalamus during photoperiodism and adult neurogenesis. We conclude that ^1^H-MRS could be a promising non-invasive translational technique to investigate the existence of adult neurogenesis in-vivo in gyrencephalic brains.

## INTRODUCTION

The hypothalamus (HYP) is a small structure belonging to the limbic system and situated at the base of the brain near the pituitary gland. HYP makes the link between the endocrine and the central nervous system and is responsible for the maintenance of the body’s internal balance. It plays a crucial role in several functions such as the regulation of appetite and body temperature, the control of reproduction and sexual behavior, the release of hormones, the control of daily physiological cycles and of emotional responses (1). This important structure is densely populated with cells that are able to regulate glucose (Glc) availability by sensing various metabolites (glutamate (Glu) and γ-aminobutyric acid (GABA)) to regulate energy balance, among which astrocytes, tanycytes and microglia (2).

Metabolic investigations within various specific structures of HYP in different species have been abundantly performed *in-vitro* (3, 4). More recently, neuroimaging (Positron Emission Tomography (PET) and magnetic resonance imaging (MRI)) studies both in rodents and in the human contributed to the *in-vivo* evaluation of the relationships between altered feeding behavior or reproductive activity and hypothalamic responses (5) as well as the evaluation of functional connectivity between the hypothalamic structures and other brain areas (6). These investigations led to a better knowledge of hyperemic mechanisms, dopaminergic circuits, reward-based theories, Gonadotropin-Releasing Hormone (GnRH) neuronal system, etc… within hypothalamic nuclei, which may also provide new ways for the evaluation of risk factors for eating and reproductive disorders and for the design of prevention strategies. Despite its small size and depth within the brain, non-invasive approaches such as proton MR spectroscopy (^1^H MRS) were conducted successfully in the HYP of rodents and humans to explore normal and altered hypothalamic metabolism (7, 8, 9, 10). Notwithstanding these pioneering investigations, HYP remains an underexplored structure of the brain and novel strategies are needed for a more accurate understanding of the mechanisms underlying alterations of the functions it controls (11).

Adult neurogenesis (AN) is the process of continuously generating new cells (neurons and glial cells) that can functionally integrate pre-existing circuits in the mammalian brain throughout life. AN recapitulates the developmental process of embryonic neurogenesis from proliferation to differentiation, migration, axonal and dendritic development and finally synaptic integration of new born neurons. This process occurs in specific areas of the brain, namely the sub-granular zone (SGZ) of the dentate gyrus, the subventricular area lining the lateral ventricles (SVZ) and the tuberal hypothalamus (12, 13, 14, 15, 16, 17).

The hypothalamic neurogenic niche has driven less attention compared to the other neurogenic niches but its implication in metabolic, eating disorders and reproductive functions represents a major area of research (18, 19,20). While structural and functional aspects of AN have already largely been investigated, little is known about metabolic regulation of AN although recent studies suggest that the metabolism ensures life-long addition of new neurons in the mammalian brain (21).

Sheep belong to a large and long-lived mammalian species with long development and life expectancy and possess a gyrencephalic brain. Studies of the hippocampal, olfactory and hypothalamic AN in sheep revealed distinctive features in the dynamics of neuronal maturation compared to rodents and appears to exhibit longer maturation time for new neurons (22, 23). AN of the sheep HYP is strongly modulated by photoperiod (17, 24). This physiological response of organisms to the ratio of the day to the night length represents a developmental strategy of conservation, having a direct impact on reproduction by alternating periods of sexual rest and sexual activity. High levels of hypothalamic cellular proliferation (17, 22, 24) are associated to short photoperiod (short days (SD), from September till January) whereas basal levels of hypothalamic cellular proliferation are observed during long photoperiod (long days (LD), from April till July). We believe that the investigation of a potential metabolic link between photoperiod and AN using non-invasively acquired biomarkers such as metabolite concentrations could bring new insights to the role of AN. Moreover, the sheep model used in the present study may represent a promising alternative animal model for translational purposes.

In the present work, ^1^H-MRS was conducted longitudinally in the HYP of adult ewes at 3T to investigate metabolic changes due to the photoperiod. The present paper is a part of a more extended work that included eight ewes and a different multiparametric MRI protocol.

## MATERIALS AND METHODS

### Animals

This study was approved by the Val de Loire animal experimentation ethics committee (CEEAVdL) in accordance with the guidelines of the French Ministry of Agriculture for animal experimentation and European regulations on animal experimentation. All experiments conform with the ARRIVE guidelines and were performed in accordance with regulations regarding animals (authorization N° 22353 of the French Ministry of Agriculture in accordance with EEC directive). All experiments were performed in 8 adult Ile-de-France (IF) ewes aged 3 to 4-years old, gathered under standard husbandry at the INRAE Val-de-Loire research center (Nouzilly, Indre-et-Loire, France, 47°33′00.8”N 0°46′55.3”E). The IF breed shows seasonal reproductive cycles that are well characterized (25, 26). The experimental facilities are approved by the local authority (agreement number E37–175–2). All animals were fed daily with hay and pellets and had *ad-libitum* access to water.

Animals were examined at four time points (P01, P02, P03 and P04) from May (P01) till July (P04) during long days (LD) and at four time points from late September (P01) till November (P04) during short days (SD). These time points during LD and SD were chosen since they correspond to basal and increased levels of AN respectively (16, 17). Due to time constraints, 4 ewes underwent ^1^H-MRS under protocol P1 while the 4 remaining ewes underwent the functional MRI protocol P2 as outlined in Fig.1.. Time points were spaced two weeks apart to allow for recovery from anaesthesia. Blood samplings twice per week were performed from March till November to follow each ewe’s hormonal status by measuring plasmatic progesterone concentration (Fig.1). Briefly, after catheterization of the jugular vein, approximately 5 ml of blood from the jugular vein and always at the same time of day (9:00 h) were drawn in vacutainers. After centrifugation, blood plasma was transferred into smaller tubes and deep-frozen. At the end of November, once all the blood samples had been collected, plasma progesterone was assayed by the radioimmunoassay method described by Terqui and Thimonier (27). When progesterone concentration was lower than 0.75 ng·mL^−1^ of plasma, the ewe was considered to be in the follicular phase of the cycle or in anovulation. As expected, from September to December, which corresponds to the breeding period, all ewes are cycling (show cyclic progesterone concentrations) (Fig S1A) indicating an increased sensitivity of the reproductive axis to short days.

**Figure 1:**
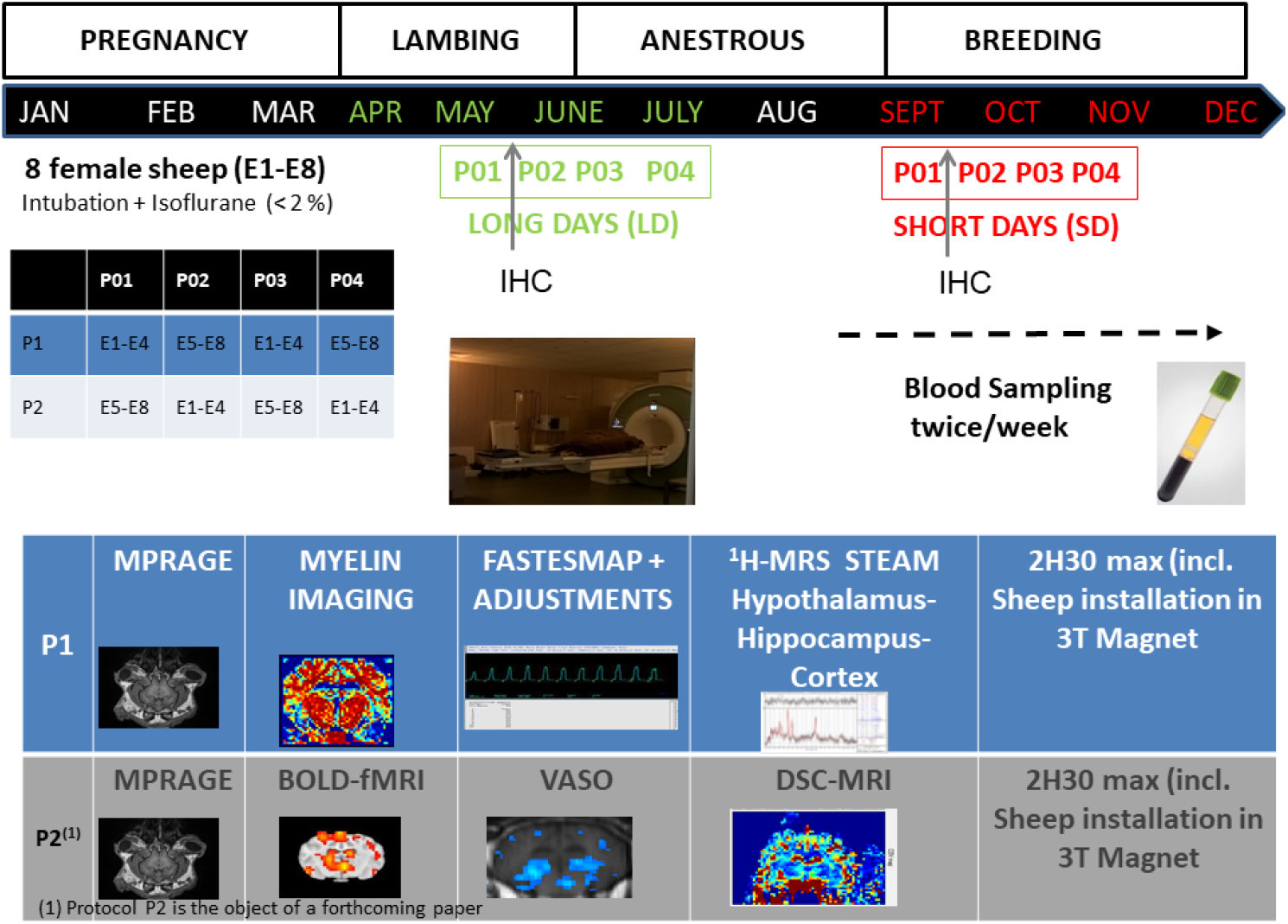
Timeline and Protocols: 8 female sheep underwent MRI and MRS at 3T following 2 interleaved protocols (P1 and P2) as described in the table displaying the time points (P01, P02, P03, P04) and the protocols (P1, P2) for MR scans of each ewe for an enhanced sensitivity of structural, functional and metabolic parameters to the photoperiod and the adult neurogenesis (AN) within the hypothalamus. Photoperiod describes the sensitivity of the sheep brain to the ratio of day to night length. Therefore, the sheep were scanned from May till July during long days (LD) and from late September till November during short Days (SD), which also corresponds to the period of sexual activity in sheep and the period of increased AN. During LD and SD, MRI and MRS took place at 4 time points (P01, P02, P03 and P04). During SD, blood samplings took place to determine the physiological status of the ewes (luteal or follicular phase of the cycle). 4 ewes were scanned per time point according to protocols P1 and P2. Protocol P1 involved structural, myelin imaging and ^1^H-MRS of the hypothalamus while Protocol P2 involved structural, functional and perfusion imaging. The duration of both protocols was 2H30. Myelin imaging and protocol P2 are the object of another publication currently under writing. Sacrifice and immunohistochemistry (IHC) were performed in a different cohort of ewes between P01 and P02

The day before MRI experiments, 3 ewes were transported to the MR facility. Care was taken to always provide company to individual animals to avoid stress since sheep are a gregarious species. Each animal was fasted 24-hours prior to intubation. Following immobilization, the sheep was intubated after intravenous administration of a mixture of ketamine and xylazine (20 mg/kg). Each ewe was transported to the MRI room, installed prone on the MRI bed (Fig 1) and anaesthesia was immediately switched to 1.5-2% isoflurane (28) in medical air through a respirator (Aestiva, GE Healthcare, Datex-Ohmeda, USA). The respirator allowed continuous control of respiration rates. An oximeter was attached to one of the hind-paws allowing for the control of the partial pressure of oxygen and heart rate. The temperature was controlled through MRI-compatible rectal probe. The duration of each MRI session was 150 minutes for each animal. No rumenostomy was performed. Frequent checks by a veterinary were performed during experiments. Animals were removed from the MRI when intra-abdominal pressure was estimated to be too high. All ewes were scanned between 8 am and 3 pm over 3 days per time point under the exact same conditions.

### Magnetic Resonance acquisitions

MR imaging was conducted on a 3T whole body MR Scanner (Siemens Verio, Erlangen, Germany) with a large 4 channel flex coil surrounding the entire head. Following the acquisition of pilot images, T_2_-weighted multislice images were acquired covering the entire sheep brain for further slice positioning. Structural images were acquired using the T_1_-weighted 3D magnetization prepared rapid gradient echo (MPRAGE) sequence (TR/TE/TI=2500/318/900 ms; Flip angle = 12; NEX= 2; FOV=192 × 192 mm^2^, Matrix size = 384 × 384; Voxel size = 0.5 × 0.5 × 0.5 mm^3^).

Proton MRS (^1^H-MRS) was conducted using a STEAM sequence (2048 points and a bandwidth of 3000 Hz) and the following parameters (TR/TE/TM= 3000/5/30 ms; 256 acquisitions) in a large voxel of interest (VOI = 10 × 12 × 13 mm3) covering the entire HYP. FASTESTMAP (29) was used for shimming down to a water linewidth of 10 ± 2Hz within the hypothalamus. Both techniques came within the MRS package developed by Edward J. Auerbach and Małgorzata Marjańska and provided by the Center for Magnetic Resonance Research (CMRR) of the University of Minnesota under a C2P agreement. Collected MRS data were analyzed using LCModel (30) and metabolite quantification was obtained using the unsuppressed water signal acquired in the same VOI (16 acquisitions).. The water suppression was performed using a VAriable Power and Optimized Relaxations delays (VAPOR) module (31). Neurochemicals quantified with Cramér-Rao lower bounds (CRLB) to estimate the errors of the neurochemical quantification under 40 % were considered reliable.

The basis set was provided by Stephen Provencher (Stephen Provencher Inc., Oakville, Ontario, Canada), included simulated macromolecules and incorporated the following metabolites: alanine (Ala), Aspartate (Asp);Cr (Creatine); γ-amino butyric acid (GABA), Glucose (Glc), Phosphocholine (PCho), Glycerophosphocholine (GPC), Glutamate (Glu), Glutamine (Gln), N-Acetyl-Aspartate (NAA), *N*-Acetylaspartylglutamic acid (NAAG), Glucose (Glc), Lactate (Lac), Scyllo-Inositol (Scyllo), Taurine (Tau), myo-Inositol (mI). Moreover, the following sums were also outputs of LCModel:

Total NAA (tNAA = NAA + NAAG), Glutamine + Glutamate (Glx), Total Choline (tCho).

### MRS voxel tracking and tissue segmentation

the MRS VOI position was tracked using a 2D T1-weighted Spin –Echo sequence (TR/TE=500/8.4 ms; FOV=220 × 200 mm^2^; Matrix=128 × 128) acquired prior to MRS in the sagittal, coronal and axial planes. These images helped us for the accurate and identical positioning of the VOI for each ewe.

MP2RAGE images acquired in each ewe during both LD and SD periods were segmented using Diffeomorphic Anatomical Registration Through Exponentiated Lie Algebra (DARTEL) (32) and our in-house made anatomical sheep template (33). Tissue composition inside the VOI was calculated based on the segmentation of 3D MPRAGE images using Gannet3.0 (34). Water concentrations, used in LCModel analysis, were calculated based on the volume fractions of white matter (WM), grey matter (GM) and cerebrospinal fluid (CSF), assuming water concentrations of WM, GM and CSF of 35,880, 43,300 and 55,556 mM, respectively. Metabolite concentrations were then divided by the fraction of WM and GM to correct for CSF inside the VOI, since metabolites are mainly present in WM and GM (35). The signal-to-noise ratio (SNR) was obtained using N-acetylaspartate (NAA) peak height at 2.01 ppm divided by standard deviation (SD) of noise collected betwee 8 and 10 ppm. For all spectra, LCModel quantification was performed on a spectral window between 0.2 and 4.2 ppm.

### Statistical Analysis

Data are presented as mean ± standard deviation unless otherwise stated and were analysed using the analysis of variance (ANOVA) for all metabolites together (repeated measures) followed by Bonferroni tests for post-hoc comparisons. The factors analysed were photoperiod (LD and SD) and Time (P01, P02, P03 and P04) during each period. Significance was considered for p < 0.05. Normality tests were performed on data sets using a Kolmogorov-Smirnov test. Statistical analyses were performed using GraphPad (Prism, San Diego, CA, USA).

Student t-tests were used to compare mean signal to noise (SNR) ratios and mean N-Acetyl Aspartate (NAA) to Creatine (Cr) ratios. To compare the mean normalized concentrations between LD and SD, a Mann-Whitney non parametric 2 tailed test was performed.

### Immunohistochemistry

#### Animals

Île-de-France primiparous non-gestating ewes were used in this study (n=10, 1.75±0.25-year-old). Animals were kept in sheepfold, under natural photoperiods, fed daily with alfalfa, corn, straw and a supplement of vitamins and minerals and had *ad libitum* water. Animals were handled and cared for in accordance with the EC Directive of 24 November 1986 (86/609/EEC) and with authorization A37110 from the French Ministry of Agriculture.

#### Tissue preparation

Five brains were collected in October and five in May corresponding respectively to the short photoperiod (Short Days/SD) and the long photoperiod (Long Days/LD). Prior to fixation, brains were perfused through the carotid arteries with 1% sodium nitrite saline solution at 37°C (2l, 0.9% NaCl) to wash the vascular bed. Brains were then fixed with 4% paraformaldehyde solution at 4°C (0.1M phosphate buffer; pH 7.4). Hypothalami were dissected from the optic chiasma to the premammillary recess from fixed tissues and post-fixed in the 4% paraformaldehyde fixative solution for 48h. Hypothalamic blocks were then immersed in a 20% sucrose solution at 4°C for cryoprotection. Prior to microcutting, blocks were frozen by immersion into isopentane at −50°C. Coronal sections (14µm thick) were collected from a distance of 2.5mm rostral to the premammillary recess using a cryostat (Leica CM 3050 S). The sections were directly mounted on Superfrost Plus slides (Fisher Scientific, Illkirch, France) and stored at −80°C until used for immunohistochemistry. Three sections of the ARH were selected for immunolabeling (Fig. 6A).

#### Immunolabeling

For brain section immunolabeling, an unmasking step was included in which sections were subjected to heat mediated antigen retrieval using citric acid (pH 6). Sections were warmed to 90°C in a microwave oven in citrate buffer (0.01M, pH 6.0). After cooling to room temperature, sections were rinsed three times in TBS (0.01M, pH 7.4). For all antibodies, sections were blocked and permeabilized with TBS supplemented with 0.3% Triton and 5% mare serum respectively for 30min. The sections were incubated with primary antibodies diluted in the same solution overnight at 37°C, followed by secondary antibodies at 37°C, in the dark, for 2h (Table 3). Then, the sections were counterstained in a solution of Hoechst (1µg/1ml dilution) at room temperature for 2 min. The sections were next coverslipped using Fluoromount (Southern Biotech). For all antibodies, normal serum IgGs of appropriate species were used as negative controls.

#### Quantification

The densities of neural stem cells [SOX2+] and astrocyte [SOX2+S100+] and oligodendrocyte [SOX2+OLIG2+] progenitors in the ARH were estimated using a semi-automated analysis process, running on ImageJ (version 1.52, NIH, US) and Imaris software (version 9.6.0, Bitplane, UK) (Fig. 7B). A first step consisting in the application of a 3×3pixels median filter has been used to improve signal/noise ratio. Densities of astrocyte and oligodendrocyte progenitors were conducted within an area of interest of 800×800µm that lined the 3^rd^ ventricle, called periventricular space. [SOX2+], [SOX2+S100+] and [SOX2+OLIG2+] cells were automatically detected, using the Imaris “spot” function following the use an established grey level threshold and an estimated cell radius of 6µm. Semi-automated Hoechst-positive cell detections were used as controls.

#### Image capture

All images for figure preparation were acquired using a microscope slide scanner (Zeiss, Axio.Scan. Z1) with Zen 2.3 software (version 4.0.3, Carl Zeiss, Oberkochen, Germany). Images shown in the different figures were pseudocolored using Imaris (version 9.6.0, Bitplane, UK) and adjusted for brightness and contrast.

#### Statistical analysis

Due to the small sample size, data are presented as mean values ± Standard Error of the Mean (SEM). Statistical analyses were performed with GraphPad Prism (version 8.4.3) software. The normality of data were tested (Shapiro tests). As the data from [SOX2+] cells did not pass the normality test, comparison between seasons was performed by a Mann-Whitney test, unlike the data of [SOX2+S100+] and [SOX2+OLIG2+] cells which pass the normality test and for which the comparison between seasons was performed by an unpaired t-test. Results are presented as barplots. Differences were considered statistically significant for a p-value ≤ 0.05.

## RESULTS

### Neurochemical profiling within the hypothalamus

Figure 2 depicts T_1_-weighted MPRAGE images with the position of the VOI within the HYP in the coronal (A), sagittal and axial (C) planes.. Representative examples of raw labeled proton MR spectra acquired within these VOIs during LD (D) and SD (E) are also presented together with LCModel fitted MR spectra.. Mean SNR values were 27 ± 9 in HYP The water linewidth was 12 ± 2 Hz on average and the linewidth of total creatine was 11± 2 Hz for spectra acquired during LD and 9 ± 2 Hz for spectra acquired during SD with no significant difference (p>0.05). Examples of MR spectra obtained in the hypothalamus of ewes at each time point during LD (blue color) and their respective spectra obtained at the same time point for the same ewe during SD (red color) are shown (Fig 3). In a few examples, extracranial lipid contaminations were seen (Fig. 3). The concentrations of metabolites for Glu, Gln, NAA and mI at each time point during LD and SD and for each ewe are reported in Table S1. The neurochemical profiles within the HYP acquired at 4 time-points during LD and SD were compared (Fig. 4, metabolite concentrations ± SD). A significant effect of Time was found (F (7, 312) =6.86; p< 0.0001) while the effect of the photoperiod (or the comparison between LD and SD) was significant at P01 and P04 (F (1, 26) = 22.88; p<0.0001). Significant changes between metabolite concentrations during LD and SD were assessed using Bonferroni post-hoc tests and are summarized in Table 1. Glu (p< 0.05), mI (p<0.05), and Glx (Glu+ Gln) (p< 0.001) concentrations were significantly higher during LD compared to SD at P01. During SD, Glu, NAA and Glx concentrations demonstrated tendencies to increase at subsequent time-points and levels became significantly higher compared to LD only at P04 for Glu (p<0.05) and Glx (p<0.001) and at P04 for NAA (p< 0.05) (Fig.4)..

**Table 1.** Cramer/Rao Lower Bounds (CRLBs) ± standard deviation (sd)

| | P01-LD | ±sd | P01-SD | ±sd | P02-LD | ±sd | P02-SD | ±sd | P03-LD | ±sd | P03-SD | ±sd | P04-LD | ±sd | P04-SD | ±sd |
| --- | --- | --- | --- | --- | --- | --- | --- | --- | --- | --- | --- | --- | --- | --- | --- | --- |
| Cr | 7.50% | 0.015 | 5.25% | 0.004 | 6.75% | 0.008 | 7.33% | 0.005 | 9.00% | 0.028 | 6% | 0.01 | 7.17% | 0.01 | 6.50% | 0.01 |
| Gln | 27.50% | 0.055 | 33.25% | 0.150 | 25.50% | 0.068 | 38.67% | 0.187 | 31.00% | 0.099 | 29% | 0.05 | 34.83% | 0.19 | 23.75% | 0.03 |
| Glu | 15.50% | 0.045 | 17.75% | 0.048 | 20.75% | 0.068 | 18.33% | 0.025 | 14.00% | 0.014 | 15% | 0.02 | 19.33% | 0.04 | 12.75% | 0.02 |
| GPC | 5.00% | 0 | 3.25% | 0.004 | 4.25% | 0.004 | 8.67% | 0.045 | 5.00% | 0.014 | 8% | 0.07 | 4.33% | 0.01 | 6.00% | 0.03 |
| ml | 5.00% | 0 | 4.00% | 0.007 | 5.00% | 0.007 | 6.00% | 0.000 | 6.00% | 0.014 | 5% | 0.01 | 5.33% | 0.01 | 4.75% | 0.00 |
| NAA | 10.00% | 0.04 | 9.00% | 0.007 | 13.00% | 0.043 | 13.67% | 0.053 | 9.00% | 0.028 | 8% | 0.02 | 11.50% | 0.05 | 9.50% | 0.01 |
| GPC+PCho | 5.00% | 0 | 3.25% | 0.004 | 4.25% | 0.004 | 5.33% | 0.005 | 5.00% | 0.014 | 4% | 0.00 | 4.33% | 0.01 | 4.00% | 0.00 |
| NAA+NAAG | 6.50% | 0.005 | 5.25% | 0.004 | 7.00% | 0.016 | 8.33% | 0.012 | 9.00% | 0.028 | 6% | 0.01 | 8.50% | 0.02 | 6.50% | 0.01 |
| Glu+Gln | 11.00% | 0.03 | 14.75% | 0.033 | 13.25% | 0.038 | 14.33% | 0.033 | 11.50% | 0.021 | 11% | 0.01 | 15.50% | 0.04 | 9.75% | 0.01 |

**Figure 2:**
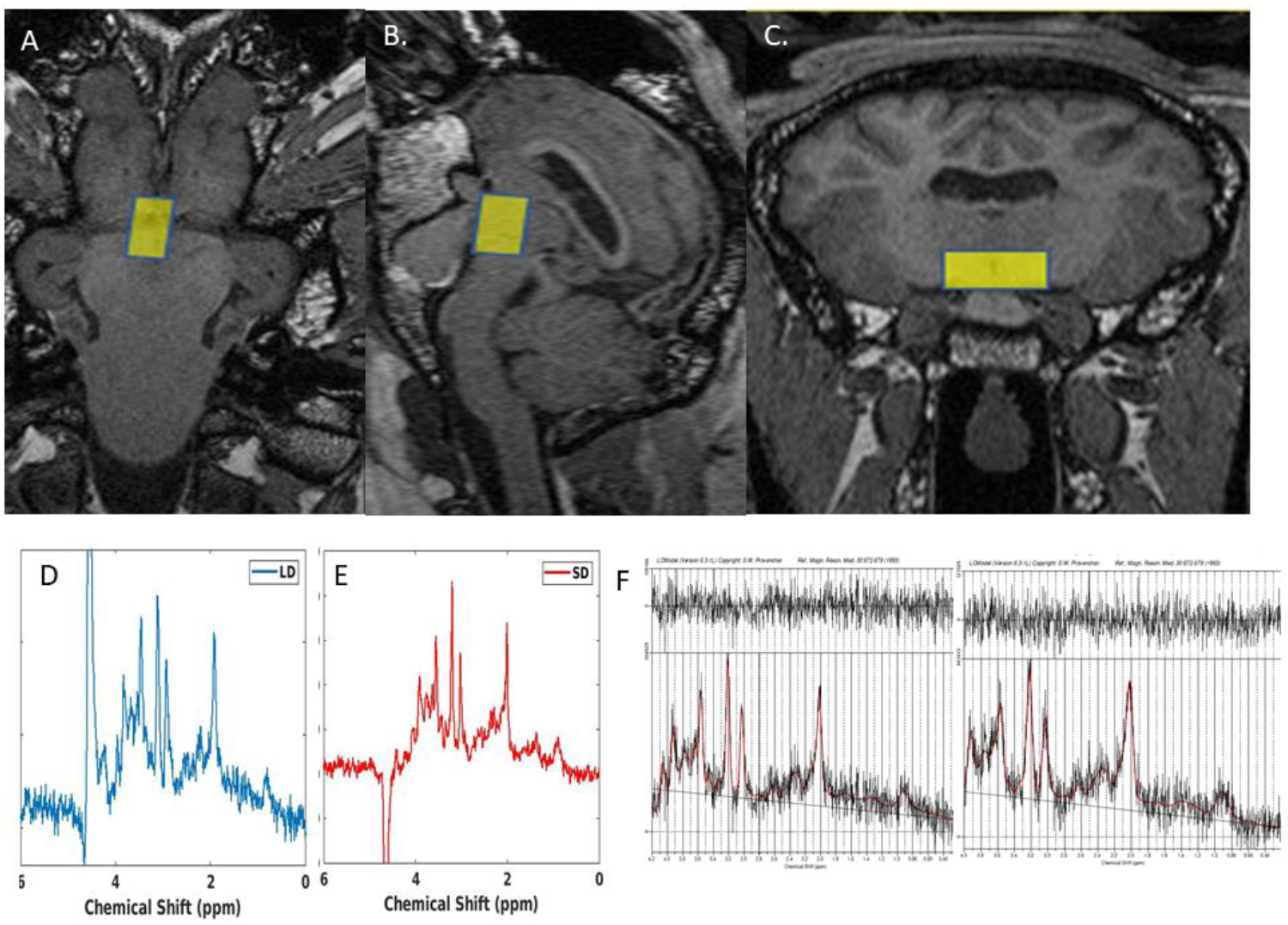
Proton Magnetic Resonance Spectroscopy (^1^H-MRS) in the hypothalamus. **A-C:** MPRAGE coronal, sagittal and axial T1-weighted images depicting the position of the VOI within the hypothalamus (VOI = 10 × 12 × 13 mm3) and **D-E:** the corresponding MR spectra acquired during LD and SD F. Representative examples of LCModel fit to a MR spectrum within the hypothalamus of a ewe.

**Figure 3:**
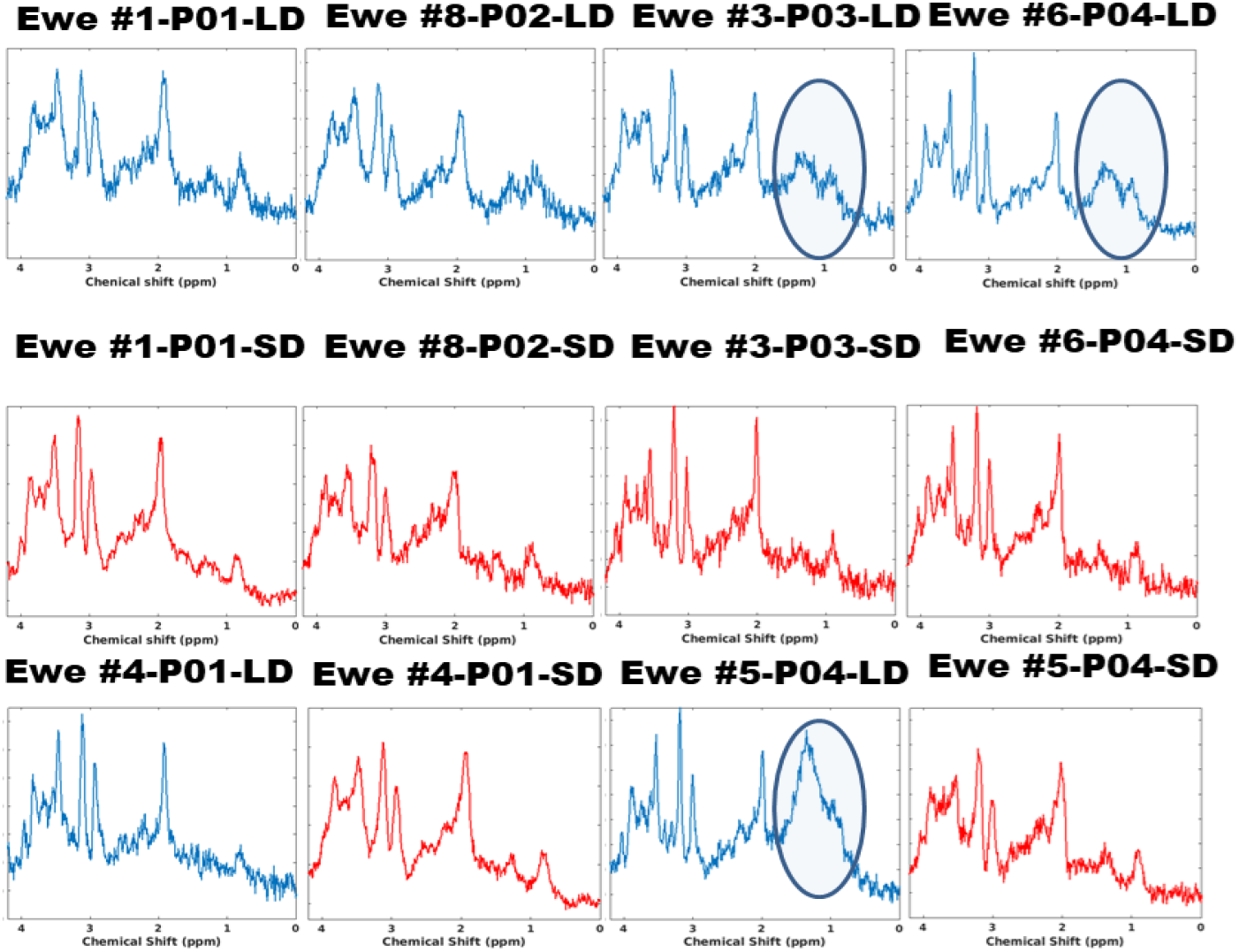
Examples of MR spectra acquired at each time points during LD and SD for different ewes. Spectra are shown for the same ewe during LD (blue) and during (SD) (red). Free-Induction decays (FIDs) were Fourier transformed and 2 Hz line broadening as well as zero and first order phasing corrections were applied to each spectrum. Unfortunately, some spectra were positioned too close to extracranial boundaries and lipid contaminations were detected as delineated in circles.

**Figure 4:**
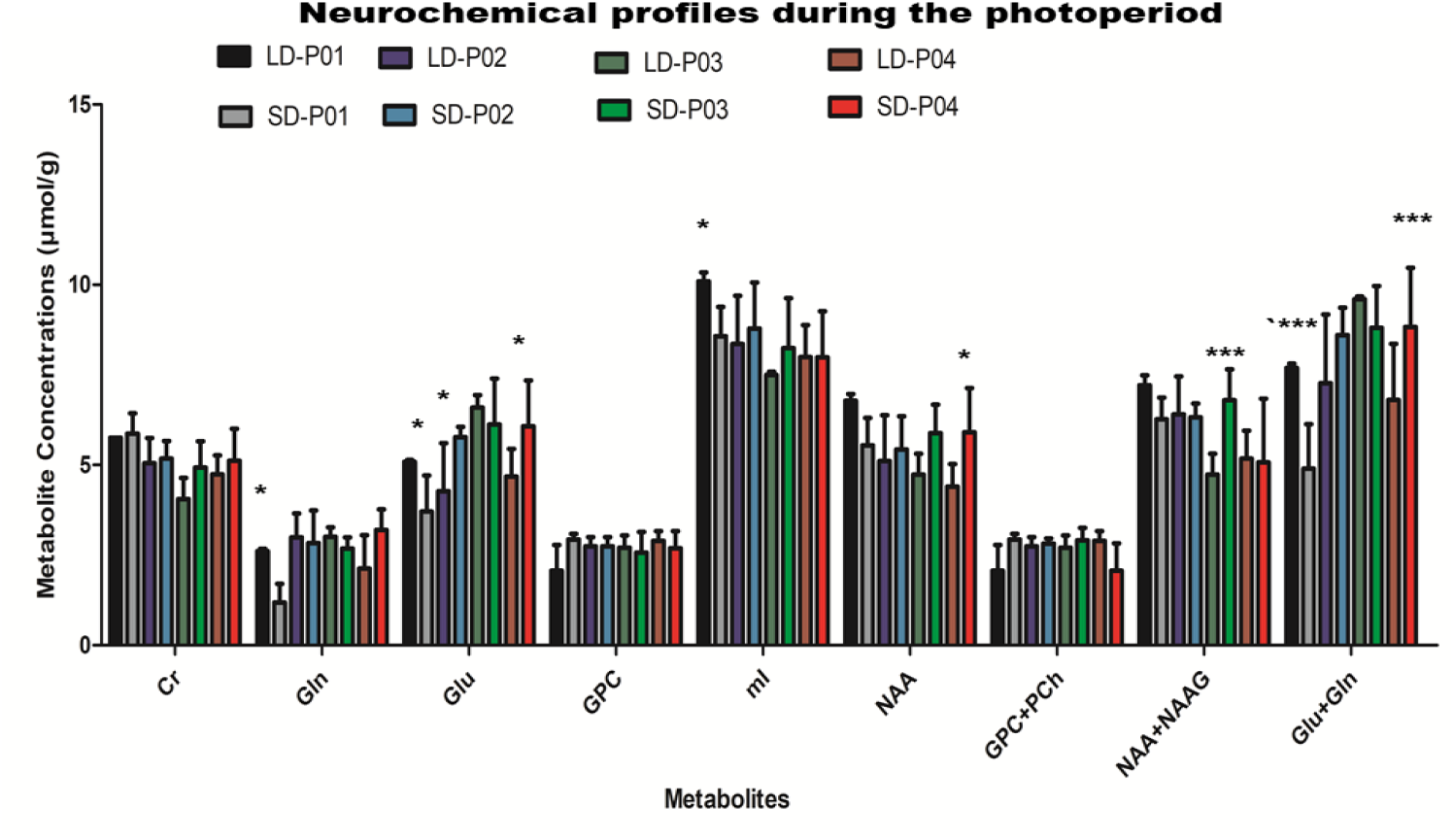
Neurochemical profiles within the hypothalamus: Neurochemical profiles of the sheep hypothalamus at each time point during LD and SD respectively (Concentrations ±SD). The statistical tests (reported in Table 2) revealed significant differences between LD and SD at P01 and P04 and between time points during LD and SD, notably for Glu, NAA, tNAA, mI and Glx. For the purpose of clarity, only significant differences between corresponding LD and SD timepoints are displayed. A 2-way ANOVA and Bonferroni correction was conducted * p<0.05; ** p<0.01; *** p<0.001

Interestingly, the number of metabolites undergoing changes per time point was more important during LD than SD, which was unexpected. mI concentrations were significantly more elevated at P01 compared to all the other points during LD (See Table 2). During SD, mI concentrations were also significantly correlated to progesterone levels (r=0.62, p0015), which was not the case for Glu (Fig. S1B). NAA and total NAA (tNAA = NAA + NAAG) concentrations decreased significantly at P03 and P04 compared to P01 and P02 during LD. This effect can also be observed when comparing the ratio of NAA to Cr levels as a function of time during LD and SD (Fig. 5A). A significant difference was observed at P04 (Student t-test, p= 0.02). Glu and Glx concentrations increased significantly at P03 compared to P01 and P02 and decreased again significantly at P04 (See Table 2, Fig 4). With regard to energy expenditure, significantly lower levels of Cr were found at P03 compared to P01 during LD. For an improved understanding of metabolic changes between SD and LD, the metabolite concentrations were normalized to the metabolic concentrations measured on the first time point of each period (SD and LD), which assumes relative changes of 100 % at P01. Fig.5B-F compare the temporal evolution of normalized concentrations between SD and LD for different metabolites. In addition, the mean relative concentration changes were compared for each period (Fig. 5G-L). During SD, Glu, Gln, NAA and mI relative concentrations were enhanced as a function of time compared to LD. Normalized concentrations reached significance for Glu, Gln and NAA at P04 only (* p<0.05, 2-way-ANOVA). Mean tCho relative concentrations were smaller during SD compared to LD (p=0.04, Mann-Whitney two-tailed test) and also significantly different at P04 (* p<0.05, 2-way-ANOVA). Mean mI and Gln and Glu/Gln normalized concentrations were not different between SD and LD (p = 0.07, Mann-Whitney two-tailed test) although the Glu to Gln ratio showed a tendency to decrease during SD compared to LD suggesting a slowdown of the Glu to Gln cycling since both Glu and Gln normalized concentrations increased.

**Table 2:** 2-way ANOVAs and Bonferroni post-hoc tests were conducted to compare the metabolite concentrations between time points P01,P02,P03 and P04 during LD and SD.(NS: Non significant, * p<0.05;**p<0.01; *** p<0.001).

|  | LD-P01 | LD-P02 | LD-P03 | LD-P04 |
| --- | --- | --- | --- | --- |
| LD-P01 |  | ml**,NAA** | Cr**,Glu*,ml***,NAA***,tNAA***,Glx** | ml***,NAA***,tNAA*** |
| LD-P02 | ml**,NAA** |  | Glu***,Glx***,tNAA** | NS |
| LD-P03 | Cr**,Glu*,ml***,NAA***,tNAA***,Glx** | Glu***,tNAA**,Glx*** |  | NS |
| LD-P04 | ml***,NAA***,tNAA*** | NS | Glu**,Glx*** |  |

|  | SD-P01 | SD-P02 | SD-P03 | SD-P04 |
| --- | --- | --- | --- | --- |
| SD-P01 |  | Gln**,Glu***,Glx*** | Gln**,Glu***,Glx*** | Gln**,Glu***,Glx*** |
| SD-P02 | Gln**,Glu***,Glx*** |  | NS | NS |
| SD-P03 | Gln*,Glu***,Glx*** | NS |  | tNAA** |
| SD-P04 | Gln***,Glu***,Glx*** | NS | tNAA** |  |
| LD-P01 | Gln*,Glu*,ml*,Glx*** | NS | ml** | ml***,tNAA*** |
| LD-P02 | Glu**,Glx*** | Glu* | Glu**,Glx* | Glu**,Glx* |
| LD-P03 | Cr**,Gln**,Glu***,tNAA*,Glx*** | tNAA* | tNAA*** | NS |
| LD-P04 | Glx** | Glx** | Glu*,NAA* | Glu*,NAA*,Glx*** |

**Table3:** Primary and secondary antibodies used for immunohistochemistry

| **Antigen** | **Antibody** | **Supplier/code** | **Dilution** |
| --- | --- | --- | --- |
| SOX2 | Anti-SOX2 goat polyclonal | R&D systems (US), #AF2018 | 1:500 |
| OLIG2 | Anti-Olig2 rabbit polyclonal | Millipore (USA), #AB9610 | 1:500 |
| S100 | Anti-S100 rabbit polyclonal | DAKO (USA), #Z0311 | 1:500 |
| **Species** | **Conjugated** | **Supplier/code** | **Dilution** |
| Donkey anti-goat IgG | Alexa488 | Molecular Probes (USA), #A21202 | 1:1000 |
| Donkey anti mouse IgG | Alexa488 | Molecular Probes (USA), #A21202 | 1:1000 |
| Donkey anti rabbit IgG | Alexa 488 | Molecular Probes (USA), #A21206 | 1:1000 |

**Figure 5:**
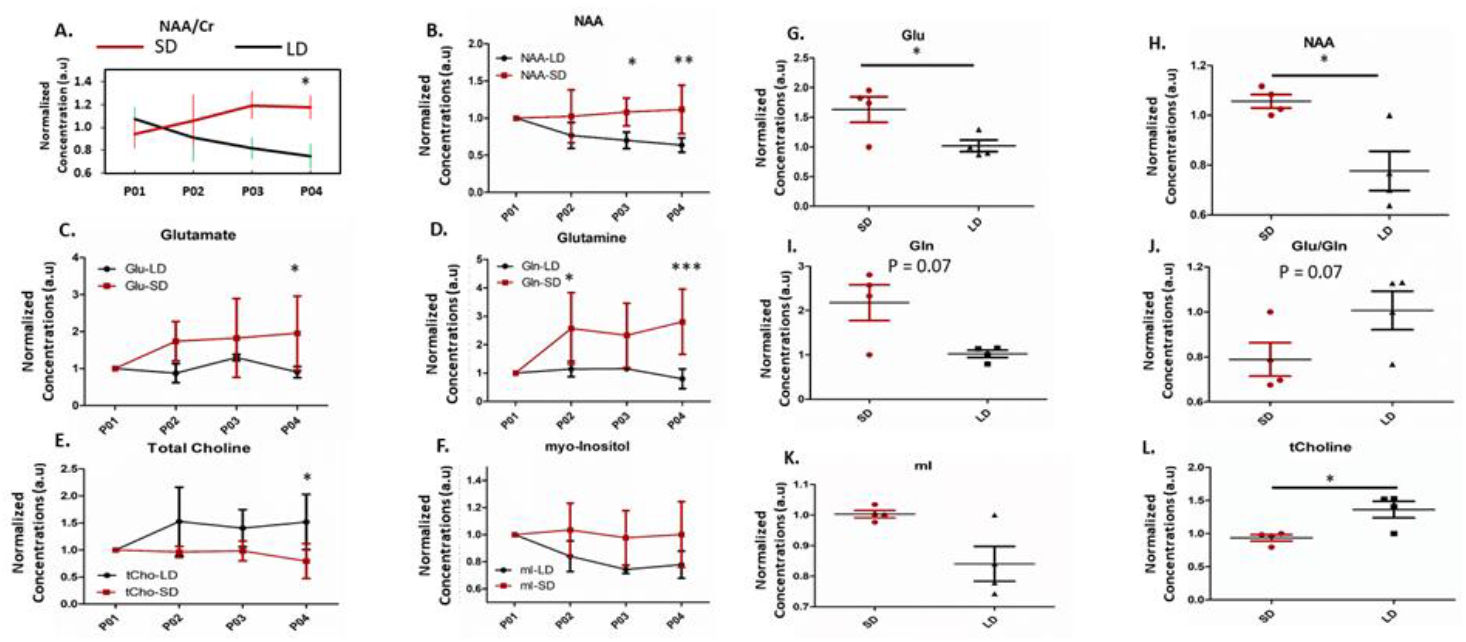
Comparison of normalized metabolite concentrations between SD and LD as a function of time (± standard deviations). **A:** Comparison of hypothalamic NAA to Cr ratio during LD and SD as a function of time. p<0.05 at P04. **B:** Normalized NAA concentrations as a function of time. p<0.05 at P03, p<0.01 at P04. **C:** Normalized Glu concentrations as a function of time. p<0.05 at P04. **D:** Normalized Gln concentrations as a function of time. p<0.05 at P02, p<0.001 at P04. **E:** Normalized tCho concentrations as a function of time. p<0.05 at P04. F: Normalized mI concentrations as a function of time. NS. **G-L:** Mean normalized Glu, NAA, Gln, Glu/Gln, mI, tCho (Mann-Whitney two tailed non-parametric test, * p= 0.02).

### Higher neural stem cells, glial and oligodendrocyte progenitor cell densities in the arcuate nucleus (ARH) during SD

To investigate whether metabolic changes that occurred according to the seasons in the hypothalamus may reflect variations in the density of cell populations, we analyzed neural stem cells ([SOX2+]), and astrocyte ([SOX2+S100+]) and oligodendrocytes ([SOX2+OLIG2+]) progenitor densities in the periventricular space of the ARH during SD and LD (Fig. 6). Using an immunohistological approach associated to a semi-automated detection (Fig. 6A) we showed an increase in [SOX2+] densities (n=10, Mann-Whitney), as well as in [SOX2+S100+] and [SOX2+OLIG2+] (n=10, unpaired t-test) cell densities during SD compared with LD (Fig. 6B-C).SD

**Figure 6:**
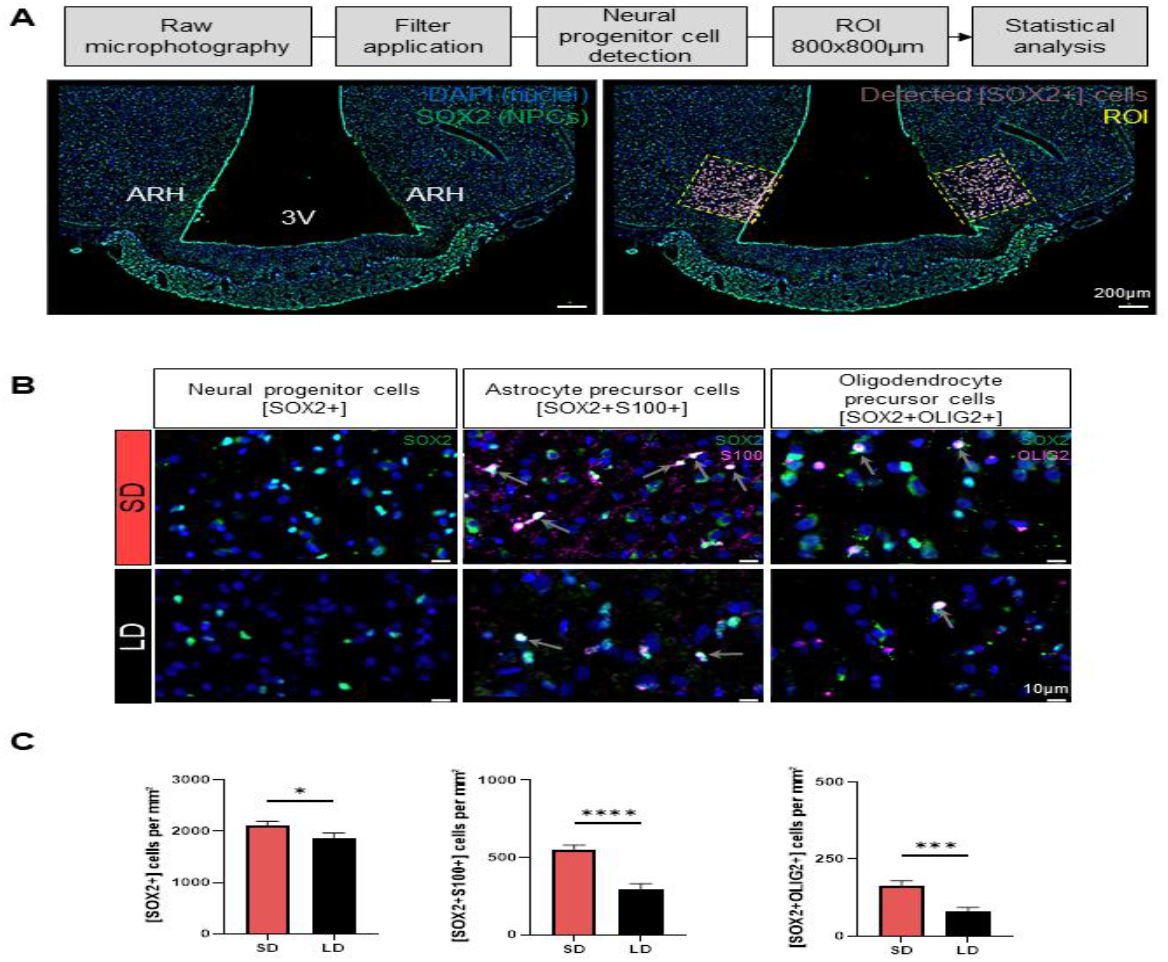
Higher densities of neural stem cells, glial and oligodendrocyte progenitor cells were found in the ARH during SP. A: Processing workflow used for the analysis of ARH cell densities. A median filter (3×3pixels) was applied on raw photomicrographs before semi-automatic detection of the markers neural stem cells [SOX2+], astrocyte [SOX2+S100+] and oligodendrocyte [SOX2+OLIG2+] progenitors respectively, performed on Imaris software. In the example, [SOX2+] are pseudocolored in green, while cell nuclei are stained with Hoechst in blue (left panel). The semi-automated detection of [SOX2+] is represented by white dots (right panel). B: Photomicrographs showing ARH neural stem cells, and astrocyte and oligodendrocyte progenitors during LD and SD. [SOX2+] labelings are in green. [S100+] and [OLIG2+] cells are in magenta. Nuclei are stained with Hoechst (blue). Grey arrows show double labelled cells ([SOX2+S100+] and [SOX2+OLIG2+]). Scale bar: 50µm. C: Panels from left to right show the mean number ± SEM of [SOX2+], [SOX2+S100+] and [SOX2+OLIG2+] respectively, estimated per mm^2^ in the ARH depending on seasons (n=5). Data are represented as barplots. For [SOX2+] a Mann-Whitney two tailed non-parametric test was used: *: p≤ 0.001. For [SOX2+S100+] and [SOX2+OLIG2+] unpaired t-tests were used: ***p ≤0.0001; **** p ≤0.0001.

## DISCUSSION

Most organisms on earth, humans included, have developed strategies to cope with environmental day-night and seasonal cycles to survive (36). For most of them, their physiological and behavioral functions, including the reproductive function, are synchronized with the annual changes of day length, to ensure winter survival and subsequent reproductive success in the following spring. In sheep, as in other seasonal mammals, several nuclei of the HYP are linked to photoperiodism with specific adaption to circadian or circannual rhythms (37). Depending on the time of the sideral year, variations in the duration of melatonin secretion either trigger or stop the breeding season. The resulting changes in hypothalamic metabolism may drive or support adult neurogenesis (37, 38). Variations in melatonin secretion duration trigger changes in hypothalamic metabolism, which may drive or support adult neurogenesis during SD in sheep (39).

In the present paper, we proposed an early exploration of the neurochemical changes induced by photoperiod in the sheep HYP. We conducted in-vivo ^1^H MRS at 3T in the HYP. To the best of our knowledge, this work represents the first characterization of the neurochemical consequences of photoperiodism with non-invasive MRS methods in the sheep brain. We will discuss our findings in relation to previous studies using invasive or ex-vivo techniques in other species and in relation to immunohistochemical data also acquired in aged-matched ewes from a different cohort to discuss the perspectives and limitations of the present study.

### The photoperiod induces significant metabolic changes within the hypothalamus

For an improved characterization of the metabolic changes, ^1^H-MRS measurements were performed longitudinally across 4 time points between April and July during LD and between September and November during SD. The time points chosen during LD and SD correspond to the basal level and the peak of AN, occurring at the mid-time of their sexual rest period and sexual activity period respectively for the IF breed (25).

Within the sheep HYP, the levels of Glu, Gln, mI and NAA changed significantly between LD and SD. Interestingly, Glu and Gln concentrations decreased by 20% (p<0.05) and 49% (p<0.05) respectively between the first time points (P01) of each period. Moreover, mean concentrations of Glu and Glx were the lowest at P01 during SD and were significantly smaller compared to all other time points (see table 2). In accordance with these findings, 31% and 48% decreases of Glu transporters (GLAST) expression in the arcuate nucleus and the tanycytes of hamsters in SD (40) were found, respectively. Moreover, the expression of Glutamine Synthetase (GS), which controls the balance between Gln and Glu in tanycytes, was also significantly decreased (40). Tanycytes lining the third ventricle constitute a large glial cell population present in the basal HYP that shares features with astrocytes and radial glial cells. Their crucial role in the maintenance of brain homeostasis, notably as glucosensors, has been outlined. In particular, their regulation of the exchanges between blood, brain and cerebro-spinal fluid (CSF) (41), during which they contribute to the exchange of metabolic information between blood and neurons or CSF and neurons. In relation with photoperiodic changes, tanycytes could be responsible for conveying regulated information between capillaries and neurons across the blood brain barrier. In Siberian Hamsters, it was suggested that tanycytes could serve as reservoirs of metabolites during SD for such transport (40). Although, no known role of metabolites released by tanycytes has been reported so far, it would represent a direction of interest for future explorations within the normal and altered metabolism of the HYP. Also, the Glu to Gln ratio relative to P01 showed a tendency to decrease during SD while both Glu and Gln normalized concentrations were higher, potentially suggesting that the Glu-Gln cycling slows down during SD due to “retention” of either Gln or Glu by tanycytes. Therefore, our data suggest the implication of the Glu-Gln cycle in the regulation of HYP metabolism according to photoperiod changes. Moreover, the increased activity of neuronal populations within the HYP during SD may require enhanced usage of Glu as initially suggested in (40) and as evidenced by the simultaneous usage of NAA at P01. Notably, NAA can act as a reservoir for Glu in case of rapid functional and structural needs during dynamic signaling demands or stress (such as survival) of neurons, astrocytes or oligodendrocytes (42).

Alternatively, early reductions of Glu, Gln and NAA concentrations at P01 could also be attributed to neuronal cell death as previously described (43). It is estimated that 30-50% of neural progenitor cells and newborn neurons undergo programmed cell death in the canonical neurogenic niches (SVZ and SGZ) within the first few weeks after birth. This process allows the elimination of superfluous cells upon regulatory mechanisms. This regulatory strategy takes place during cell differentiation and cell migration and requires utilization of both Glu and Gln in response to energetic demands. Although such selective elimination of newborn cells has not been described yet, comparative mechanisms are likely to occur for the hypothalamic neurogenic niche.

In addition, reductions in metabolic concentration are also often linked to mitochondrial functional changes. The importance of mitochondria during AN has been underlined as a key player during the differentiation and proliferation stages of AN (44). Mitochondria regulate energy metabolism and neuronal signaling in order to maintain high rates of energy demands in the brain. Imbalances in mitochondrial function lead to the disturbed development of neural stem cells during AN. This dysregulation hinders neuronal differentiation and synapse formation. During the process of AN, stem cells are subjected to switches in metabolic patterns (from high glycolytic activity to oxidative phosphorylation in terminally differentiated cells) and to oscillations in oxygen homeostasis. Episodes of hypoxia can, for example, enhance the generation of new born neurons. The Hypoxia-inducible factor 1α (HIF) is a transcription factor that allows stem cells to adapt to hypoxic environments. In particular, HIF activates genes involved in pluripotency and metabolism to sustain the functional evolution of stem cells. The analysis of HIF expression in the arcuate nucleus (ARH) by immunohistochemistry in a similar cohort of ewes to the one examined here, did not reveal a sensitivity to the photoperiod (45) despite a tendency of HIF to decrease during SD, which could be resulting from the interplay between oxygenation levels caused by AN. In support of this hypothesis, the blood oxygen level dependent functional MRI (BOLD fMRI) during a hypercapnic challenge (Protocol P2, Fig 1) (46) demonstrated no BOLD activity in the HYP during SD compared to LD at P01 (Fig S2 supplementary). By contrast, the BOLD responses to hypercapnia were augmented in other brain areas during SD.

Interestingly, the changes in Glu and Glx concentrations measured at P01 were followed by gradual increases in Glu and Glx concentrations during SD. At P04, 55 % and 39 % significant mean increases of Glu and Glx were determined respectively, during SD compared to LD. NAA (and total NAA concentrations also increased at P03 (+ 22.6%, p>0.05) and P04 (+38%, p<0.05) during SD compared to LD. At P04 during SD, the concentrations of Glu, Glx and NAA returned to the levels measured at P01 LD potentially suggesting a normalization of the HYP metabolism towards the end of the sexual activity period.

A significant decrease of mI concentrations was also demonstrated at P01 during SD compared to LD whereas levels were equivalent at subsequent time points. Moreover, mI concentrations during SD were significantly correlated to progesterone concentrations (Fig.S1B). mI concentrations are usually higher in astrocytes than in neurons and apart from being an important organic osmolyte, mI is also an intracellular messenger involved in transferring hormonal information. This role could explain the correlation pattern between mI and progesterone concentrations seen in the present study. Significant decreases in mI concentrations were also previously related to a volume-regulatory strategy of edema or to cell lysis and death further supporting our earlier hypothesis of cell death at P01 during SD (47, 48).

### Relative metabolite concentrations

We normalized the metabolite concentrations to the metabolite concentrations measured at P01 for LD and SD respectively. This normalization process allowed us to remove potential experimental or physiological differences arising from MRS measurements conducted 5 months apart in the same cohort of ewes. This corresponds to arbitrarily assuming a metabolite ratio of 100% at P01. Moreover, it highlighted the existing and almost constant metabolic differences between LD and SD, despite large standard deviations that can be attributed to the small sample size per time point.

Glu and NAA are recognized as neuronal markers whereas both mI and Gln are well-established astrocytic markers (49). The mean increases in normalized Glu and NAA concentrations were consistent with the increases in SOX2 expression, a neural stem cell marker (44), in the ARH during SD in a similar cohort of ewes examined by immunohistochemistry (Fig 6). On the other hand, normalized mI and Gln concentrations were consistent with the significant increases in the densities of glial cells and oligodendrocytes identified by SOX2/S100 and SOX2/OLIG2 co-expression respectively (Fig 6).

Finally, mean tCho was significantly decreased during SD. Choline is important for lipid metabolism in the brain and constitutes the central part of phospholipids, which are abundant in biological membranes. Choline and choline compounds are required by neural progenitor cells in the hippocampus for membrane synthesis and methylation and thereby influence neurogenesis. The significant decrease in total choline levels during SD may suggest its significant use in neurogenic processes during SD compared with LD.

### During LD, the HYP metabolic profile changed significantly

Interestingly, the number of changes in metabolite concentration was greater during LD than during SD (Table 2). LD corresponds to a period of food abundance for sheep, which remain outside in the fields at this time of the year. During LD, a continuous decrease of mI, NAA and tNAA concentrations was seen from P01 to P04, suggesting some adaption/habituation mechanisms. If these time-dependent coping mechanisms during LD were the consequence of isoflurane anesthesia and/or the time of the day at which the female sheep were scanned, the same trend should have been observed during SD, which was not the case as confirmed by the evolution in opposite directions of the NAA/Cr ratio, a neuronal marker, during LD and SD. Previous studies in rodents have shown that lower levels of NAA correlated with higher levels of dopamine release (49, 50). Dopamine is considered a neurotransmitter involved in the motivational component of the reward-motivated behaviour. The period of food abundance during LD would match with increasing levels of dopamine in the sheep brain. Moreover, dopaminergic neurons have been implicated in the transition from the breeding period to the anestrous period and vice-versa, involving an important remodelling of the hypothalamic circuitry (51), which was found intriguingly more important during the anestrous period.

Cr concentrations dropped significantly at P03 compared to P01 followed by a slight augmentation at P04. During SD a non-significant trend to Cr decrease was also observed. As herbivores, sheep do not ingest Cr through their diet and must rely on *de novo* synthesis of Cr, which might explain the observed trend. In terms of energy expenditure, it was also shown that sheep have an important resilience to hypoglycemia and can live on reserves for several days.

Finally, our immunohistochemical findings in the ARH of the HYP in another cohort of ewes (45) showed important angiogenic processes during LD, which would indeed have significant effects on the metabolism of the HYP.

### Limitations of the study

Only 4 ewes per time point underwent MRS measurements. These ewes were scanned in an interleaved manner during LD and SD allowing the observation of their HYP with both imaging and spectroscopic techniques as presented in Fig.1. Although this choice can be seen as a limitation, the ewes used in the present study were healthy and underwent MRI and MRS under identical conditions during 2 periods of the year. For ethical and practical reasons, it was not possible to scan a larger cohort of ewes. Nevertheless, the present study represents a unique opportunity to correlate multiparametric data with metabolic data (11).

Since increased AN was repeatedly observed within the HYP of the sheep during SD (24), the results were interpreted assuming a simultaneous development of new cells in this structure. However, large VOIs were used for MRS measurements although the neurogenic niche is constrained to the ARH within the HYP. Thus, metabolic observations were averaged over non-neurogenic and neurogenic tissues and cannot be considered entirely AN-dependent. Immunohistochemical data acquired in the ARH of a different cohort of ewes in our laboratory demonstrated significant neuronal, vascular (45) and glial variations in response to photoperiod (the present results). Increased spatial resolutions are therefore needed.

Here, we used a short echo STEAM sequence to acquire ^1^H MRS in the HYP at 3T. For an increased accuracy and reproducibility of metabolite quantification in such a deep brain structure, high SNR is needed. In humans, increasing the magnetic field strength from 1.5T to 3T enhanced significantly the quality of MR spectra acquired within the HYP (10). Moreover, the use of improved techniques such as short-echo semi-LASER sequences (52) enabled the longitudinal follow up of hyperglycemic patients and the preliminary assessment of a linear correlation between HYP and blood glucose concentrations (10). In rodents, MRS studies also demonstrated improved results at higher field strength (14.1 T) with more sensitive sequences, enabling the assessment of alterations of neurochemical changes in the HYP of high-fat fed and anorectic rodents (7, 8, 9). In addition, the processing of MR spectra in the sheep brain could be improved with a better tracking of motion, the measurement of sheep basis sets and of macromolecule spectra as advised in the recent consensus papers (53). Also, changes in cerebrospinal fluid partial volume were tracked (10), where CSF exchange bears important implications for a correct interpretation of photoperiod and AN-induced metabolic changes. Water concentrations, used in LCModel analysis, were calculated based on the volume fractions of WM, GM and CSF, assuming water concentrations correspond to the water concentrations for the human brain since no measurements of water content in the sheep brain was available.

Furthermore, the role of CSF in the brain and more specifically in the hypothalamus is gradually unfolding and appears significant (54) but remains difficult to understand. Notably, tanycytes, which are specialized glial cells lining the wall of the third ventricle are involved in energy homeostasis, food intake and reproduction by regulating the blood brain barrier and the blood-CSF exchanges and modulating hypothalamic neurogenesis. These cells are also affected by seasonality changes (55). Recent studies have shown that cilia, coating some of the cells lining the wall of the 3^rd^ ventricle drive CSF flow and might have significant effects on the directional CSF flow in the adjacent hypothalamus parenchyma (56). In particular, cilia-driven streams of signaling molecules offer an interesting perspective to understand better the underpinnings of CSF-borne signals dynamically transmitted to the brain. Cerebrospinal fluid (CSF)-filled ventricles resemble dynamic reservoirs of signaling substances and nutrients that can reach neurons, glia cells and stem cells within the hypothalamus. However, CSF does not flood the hypothalamus but rather delivers important factors to specific hypothalamic nuclei, which function relate to metabolism, circadian timing or the control of stem cell proliferation and differentiation. There is a concept of “volume transmission” (57), which could explain how solutes and signaling molecules are transported.

## Conclusion

Metabolic concentrations were determined for the first time using ^1^H-MRS in the sheep HYP at two opposite time points of the year in term of day length. The present study provides a framework for testing hypothesis on the impact of metabolism on the physiological role of cellular plasticity in the adult brain. Investigations on HYP and seasonal rhythms are relevant for a better understanding of alterations of the hypothalamic-pituitary-gonadal axis during appetite and reproductive disorders that may be linked to changes in AN activity levels. Additionally, with a gyrencephalic brain and the possibility to be imaged under identical conditions as humans in clinical scanners, the sheep represents an interesting animal model for further non-invasive imaging of AN and for translational purposes (28).

## ABBREVIATIONS

(Ala): Alanine
(Asp): Aspartate
(Creatine): Cr
(GABA): γ-amino butyric acid
(Glc): Glucose
(PCho): Phosphocholine
(Glu): Glutamate
(Gln): Glutamine
(NAA): N-Acetyl-Aspartate
(NAAG): *N*-Acetylaspartylglutamic acid
(tNAA = NAA + NAAG): Total NAA
(mI): myo-Inositol
(Glx): Glutamine + Glutamate
(tCho= PCh + GPC): Total Choline
AN: Adult Neurogenesis
ANOVA: Analysis of variance
ARH: Arcuate Nucleus of the hypothalamus
BOLD –fMRI: Blood Oxygen Level dependent functional MRI
CRLB: Cramér-Rao lower bounds
CSF: Cerebro-Spinal Fluid
DSC-MRI: Dynamic Susceptibility Contrast MRI
FASTESMAP: Fast, Automatic Shim Technique using Echo-planar Signal readouT for Mapping Along Projections
GM: Grey Matter
GnRH: Gonadotropin-Releasing hormone
HIF: Hypoxia-inducible factor 1α
HYP: Hypothalamus
LASER: localization by adiabatic selective refocusing;
LCModel: Linear Combination Model
LD or LP: Long days or Long photoperiod
SD or SP: Short days or Short photoperiod
MRI: Magnetic Resonance Imaging
MRS: Magnetic Resonance Spectroscopy
MPRAGE: 3D magnetization prepared rapid gradient echo
NEX: Number of excitations
SGZ: Subgranular zone
STEAM: Stimulated Echo Acquisition Mode
SVZ: Subventricular zone
SCN: Suprachiasmatic nuclei
VASO: Vascular space Occupancy
VOI: Voxel of Interest
WM: White Matter

## Author contributions

NJ and MM designed the study. NJ conducted the MR experiments. NJ, MM, PMC analyzed and interpreted the data. MB and JPD contributed to animal experiments. MM, PMC and MB, PV and DP performed and analyzed immuno-histochemical data. PMC and MM wrote parts corresponding to immunohistochemistry. NJ, PMC and MM wrote the manuscript. All authors read, corrected and approved the final version of the manuscript.

All MR related experiments were performed within the PIXANIM imaging platform of INRAE Nouzilly. We are grateful for the preparation of animals, for the precious help from all the technical staff of the platform and for granting access to the platform.

## Acknowledgements

This study was funded by a grant from Agence Nationale de la Recherche (ANR-16-CE37-0006-01) to Martine Migaud. Nathalie Just received research funding from the Lundbeck Foundation (Experiment grant, grant nr. R370-2021-402)

## Supplementary Materials and Methods

### Blood Oxygen Level Dependent functional MRI

**BOLD fMRI** was conducted using a multi-slice single shot EPI sequence (TR/TE= 2970/30 ms; flip angle = 90°; FOV = 188 x188 mm^2^; Matrix= 72 × 72; Slice thickness = 3 mm; slices = 20). 7% CO2 was delivered through tubing directly connected to the intubation system. The paradigm of stimulation consisted of 5 cycles of 60 s OFF-60 s ON periods for a total acquisition time of 11 minutes. **I**

### Image processing

Functional MR images were processed with SPM12 software (Statistical Parametric Mapping, London, UK). After pre-processing, images were co-registered to structural MPRAGE images and normalized to the in-house developed sheep brain template (1). Finally, images were spatially smoothed using a 6 × 6 × 6 mm^3^ Gaussian kernel. The general linear model (GLM) first level analysis was conducted. BOLD responses were mapped as T-value maps overlaid onto our-in-house sheep atlas template (1). Significance of BOLD responses was evaluated at cluster level using FDR-corrected p-values set to 0.01. Second level SPM analysis was conducted to compare hypercapnia to baseline at each time point during LD and SD using a one sample t-test. A pvalue<0.05 was considered significant.

## Supplementary Figures and captions

**Figure S1.**
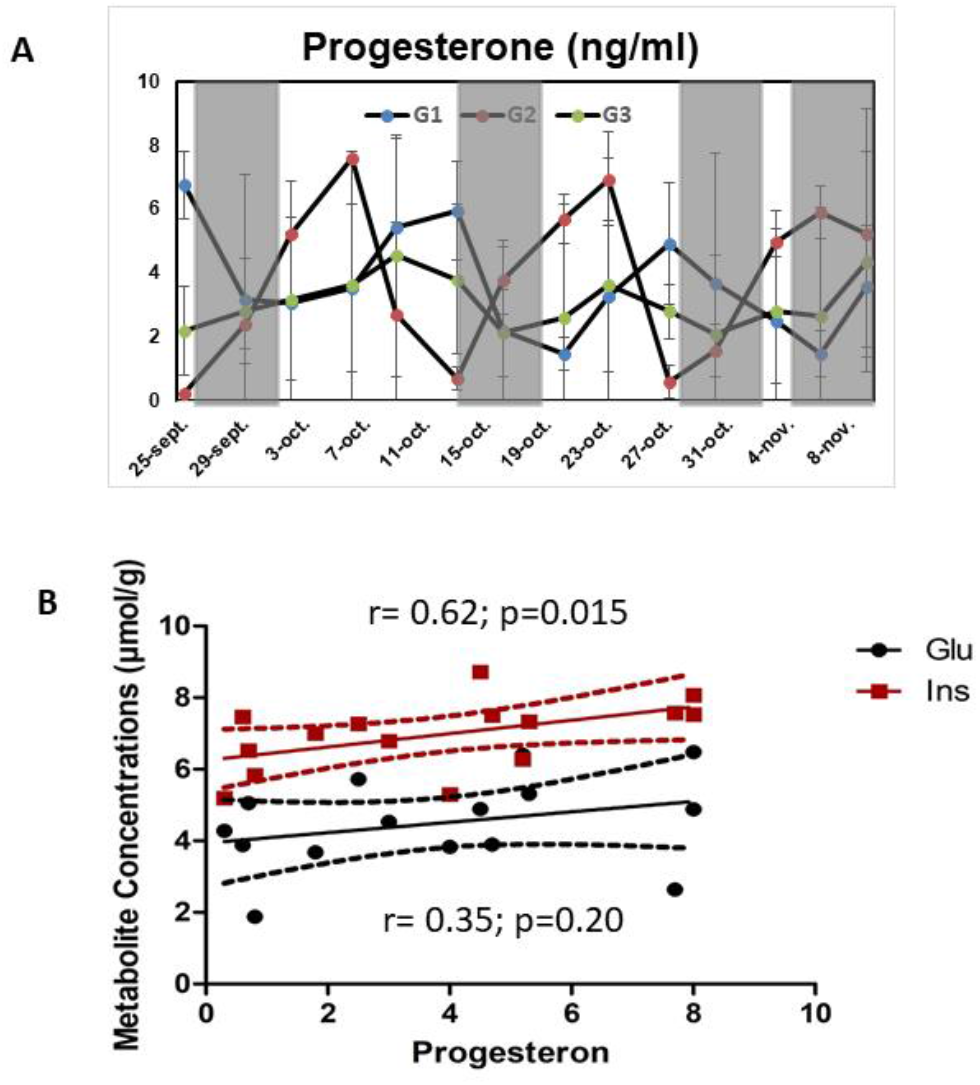
A. Cyclicity of progesterone levels during the period of sexual activity of the 8 ewes used in the present study from September to November. Shaded dark gray areas represent the MR scanning periods. Progesterone concentrations with similar dynamics (with same cyclicity) were averaged across ewes (G1 n=3 ewes, G2 n= 3 ewes and G3 n= 2 ewes) for an easier visualization. The important criterion was cyclicity and not that cycles may be shifted from one ewe to another. **B** Progesterone levels were correlated to Glu and mI concentrations during SD. Only mI concentrations were correlated to progesterone levels (r= 0.62; p=0.015).

**Figure S2:**
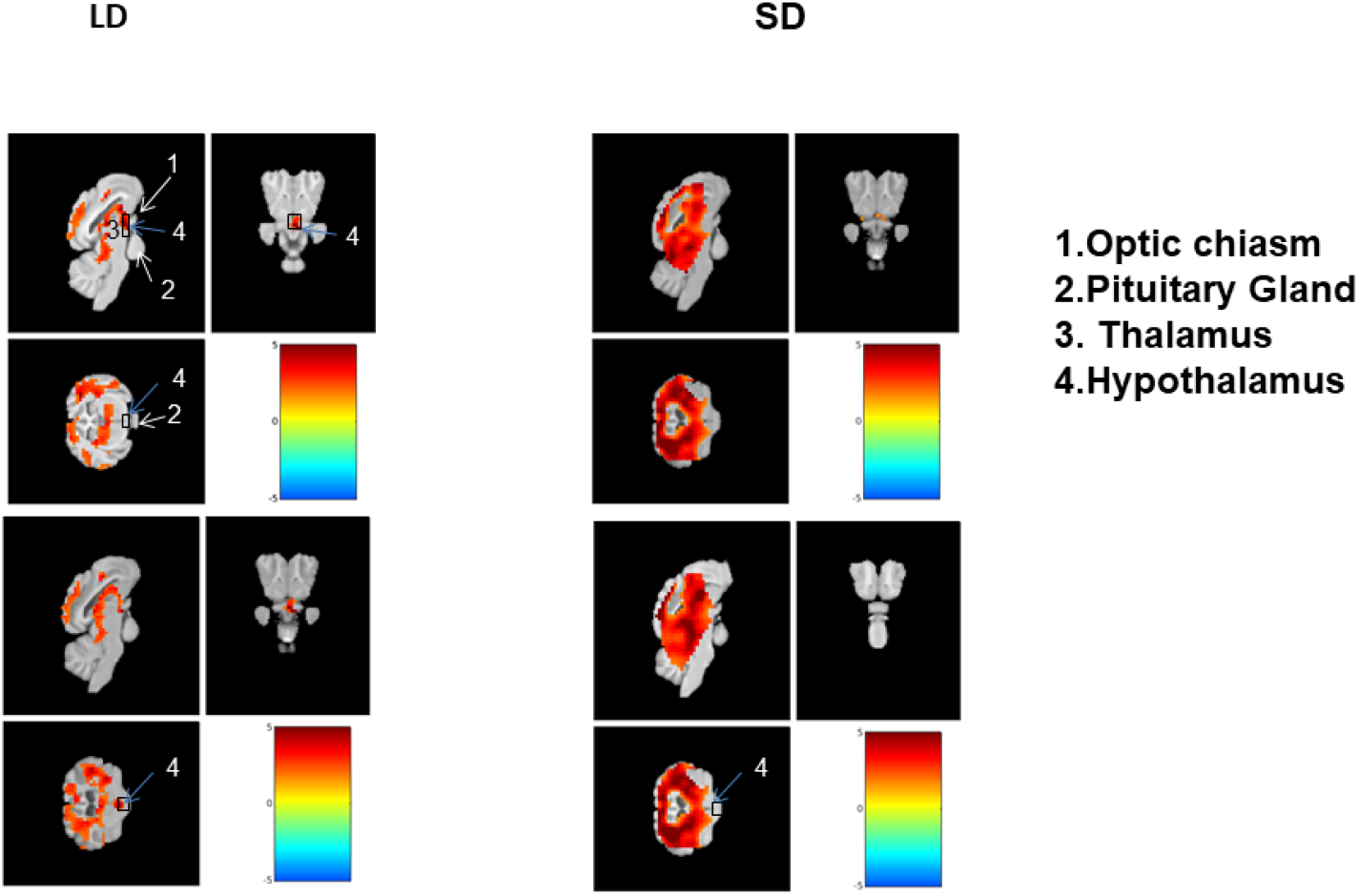
Thresholded BOLD fMRI group maps (p<0.05 Family-wise corrected) during a hypercapnic challenge within the hypothalamus of ewes (n=8) at P01 during LD and SD. No BOLD activity was detected at P01 during SD. 1. Optic chiasm. 2. Pituitary gland. 3.Thalamus.4. Hypothalamus.

**Table S1:**
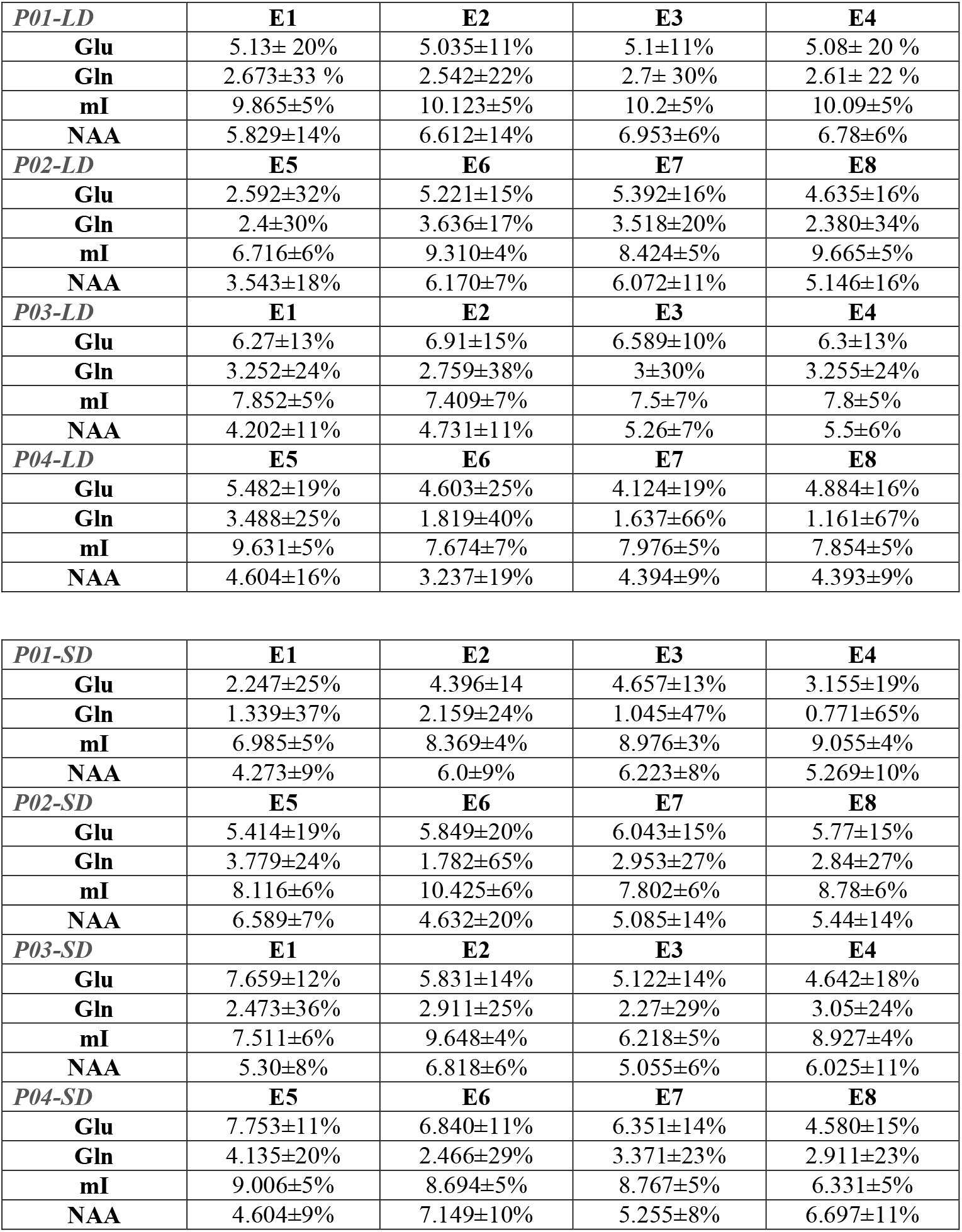
Metabolite Concentrations per ewe (E_n_) and per time point (Concentration (mM) ±CRLB (%))

| *P01-LD* | **E1** | **E2** | **E3** | **E4** |
| --- | --- | --- | --- | --- |
| **Glu** | 5.13± 20% | 5.035±11% | 5.1±11% | 5.08± 20 % |
| **Gln** | 2.673±33 % | 2.542±22% | 2.7± 30% | 2.61± 22 % |
| **mI** | 9.865±5% | 10.123±5% | 10.2±5% | 10.09±5% |
| **NAA** | 5.829±14% | 6.612±14% | 6.953±6% | 6.78±6% |
| *P02-LD* | **E5** | **E6** | **E7** | **E8** |
| **Glu** | 2.592±32% | 5.221±15% | 5.392±16% | 4.635±16% |
| **Gln** | 2.4±30% | 3.636±17% | 3.518±20% | 2.380±34% |
| **mI** | 6.716±6% | 9.310±4% | 8.424±5% | 9.665±5% |
| **NAA** | 3.543±18% | 6.170±7% | 6.072±11% | 5.146±16% |
| *P03-LD* | **E1** | **E2** | **E3** | **E4** |
| **Glu** | 6.27±13% | 6.91±15% | 6.589±10% | 6.3±13% |
| **Gln** | 3.252±24% | 2.759±38% | 3±30% | 3.255±24% |
| **mI** | 7.852±5% | 7.409±7% | 7.5±7% | 7.8±5% |
| **NAA** | 4.202±11% | 4.731±11% | 5.26±7% | 5.5±6% |
| *P04-LD* | **E5** | **E6** | **E7** | **E8** |
| **Glu** | 5.482±19% | 4.603±25% | 4.124±19% | 4.884±16% |
| **Gln** | 3.488±25% | 1.819±40% | 1.637±66% | 1.161±67% |
| **mI** | 9.631±5% | 7.674±7% | 7.976±5% | 7.854±5% |
| **NAA** | 4.604±16% | 3.237±19% | 4.394±9% | 4.393±9% |

| *P01-SD* | **E1** | **E2** | **E3** | **E4** |
| --- | --- | --- | --- | --- |
| **Glu** | 2.247±25% | 4.396±14 | 4.657±13% | 3.155±19% |
| **Gln** | 1.339±37% | 2.159±24% | 1.045±47% | 0.771±65% |
| **mI** | 6.985±5% | 8.369±4% | 8.976±3% | 9.055±4% |
| **NAA** | 4.273±9% | 6.0±9% | 6.223±8% | 5.269±10% |
| *P02-SD* | **E5** | **E6** | **E7** | **E8** |
| **Glu** | 5.414±19% | 5.849±20% | 6.043±15% | 5.77±15% |
| **Gln** | 3.779±24% | 1.782±65% | 2.953±27% | 2.84±27% |
| **mI** | 8.116±6% | 10.425±6% | 7.802±6% | 8.78±6% |
| **NAA** | 6.589±7% | 4.632±20% | 5.085±14% | 5.44±14% |
| *P03-SD* | **E1** | **E2** | **E3** | **E4** |
| **Glu** | 7.659±12% | 5.831±14% | 5.122±14% | 4.642±18% |
| **Gln** | 2.473±36% | 2.911±25% | 2.27±29% | 3.05±24% |
| **mI** | 7.511±6% | 9.648±4% | 6.218±5% | 8.927±4% |
| **NAA** | 5.30±8% | 6.818±6% | 5.055±6% | 6.025±11% |
| *P04-SD* | **E5** | **E6** | **E7** | **E8** |
| **Glu** | 7.753±11% | 6.840±11% | 6.351±14% | 4.580±15% |
| **Gln** | 4.135±20% | 2.466±29% | 3.371±23% | 2.911±23% |
| **mI** | 9.006±5% | 8.694±5% | 8.767±5% | 6.331±5% |
| **NAA** | 4.604±9% | 7.149±10% | 5.255±8% | 6.697±11% |

## References

1. Hofman MA, Swaab DF. The human hypothalamus: comparative morphometry and photoperiodic influences. Prog Brain Res. 1992; 93:133-47; discussion 148-9. doi: 10.1016/s0079-6123(08)64569-0.

2. Garcia-Caceres C, Balland E, Prevot V, Luquet S, Woods S, Koch M, Woods SC, Koch M, Horvath TL, Yi CX, Chowen JA, Verkhratsky A, Araque A, Bechmann I, Tschöp MH. Role of astrocytes, microglia, and tanycytes in brain control of systemic metabolism. Nat. Neurosci. 2019, 22, 7–14. doi: 10.1038/s41593-018-0286-y.

3. Evans JJ, Janmohamed S, Forsling ML Gonadotrophin-releasing hormone and oxytocin secretion from the hypothalamus in vitro during pro-oestrus: the effects of time of day and melatonin. Brain Res Bull. 1999, 48 (1):93–7. doi: 10.1016/s0361-9230(98)00151-8.

4. Hofmann K, Lamberz C, Piotrowitz K, Offermann N, But D, Scheller A, Al-Amoudi A, Kuerschner L. Tanycytes and a differential fatty acid metabolism in the hypothalamus. Glia. 2017, 2(2):231–249. doi: 10.1002/glia.23088.

5. Kamali A, Karbasian N, Ghazi Sherbaf F, Wilken LA, Aein A, Sair HI, Arevalo Espejo O, Rabiei P, Choi SJ, Mirbagheri S, Riascos RF, Hasan KM. Uncovering the dorsal thalamo-hypothalamic tract of the human limbic system. Neuroscience. 2020, 432:55–62. doi: 10.1016/j.neuroscience.2020.02.021.

6. Le TM, Liao DL, Ide J, Zhang S, Zhornitsky S, Wang W, Li CR. The interrelationship of body mass index with gray matter volume and resting-state functional connectivity of the hypothalamus. Int J Obes (Lond). 2020, 2(5):1097–1107. doi: 10.1038/s41366-019-0496-8.

7. Lizarbe B, Soares AF, Larsson S, Duarte JMN. Neurochemical modifications in the hippocampus, cortex and hypothalamus of mice exposed to long-term high-fat diet. Front Neurosci. 2019, 12:985. doi: 10.3389/fnins.2018.00985.

8. Just N, Cudalbu C, Lei H, Gruetter R. Effect of manganese chloride on the neurochemical profile of the rat hypothalamus. J Cereb Blood Flow Metab. 2011, 2(12):2324–33. doi: 10.1038/jcbfm.2011.92.

9. Just N, Gruetter R. Detection of neuronal activity and metabolism in a model of dehydration-induced anorexia in rats at 14.1 T using manganese-enhanced MRI and 1H MRS. NMR Biomed. 2011, 2(10):1326–36. doi: 10.1002/nbm.1694.

10. Joers JM, Deelchand DK, Kumar A, Moheet A, Seaquist E, Henry PG, Öz G. Measurement of Hypothalamic Glucose Under Euglycemia and Hyperglycemia by MRI at 3T. J Magn Reson Imaging. 2017, 2(3):681–691. doi: 10.1002/jmri.25383.

11. Just N, Chevillard PM, Migaud M. Imaging and spectroscopic methods to investigate adult neurogenesis in vivo: New models and new avenues,Frontiers in Neuroscience. 2022

12. Doetsch F, Garcia-Verdugo JM, Alvarez-Buylla A. Cellular composition and three-dimensional organization of the subventricular germinal zone in the adult mammalian brain. J Neurosci. 1997, 17: 5046–5061. doi:10.1523/JNEUROSCI.17-13-05046.1997.

13. Kempermann G. Why new neurons? Possible functions for adult hippocampal neurogenesis. J Neurosci. 2002, 2(3):635–8. doi: 10.1523/JNEUROSCI.22-03-00635.2002.

14. Lledo PM, Alonso M, Grubb MS. Adult neurogenesis and functional plasticity in neuronal circuits. Nat Rev Neurosci. 2006, 7:179–93. doi: 10.1038/nrn1867.

15. Yoo S, Blackshaw S. Regulation and function of neurogenesis in the adult mammalian hypothalamus. Prog Neurobiol. 2018, 170:53–66. doi: 0.1016/j.pneurobio.2018.04.001.

16. Migaud M, Butruille L, Batailler M. Seasonal regulation of structural plasticity and neurogenesis in the adult mammalian brain: focus on the sheep hypothalamus. Front Neuroendocrinol. 2015, 37:146–57. doi: 10.1016/j.yfrne.2014.11.004.

17. Butruille L, Batailler M, Mazur D, Prévot V, Migaud M. Seasonal reorganization of hypothalamic neurogenic niche in adult sheep. Brain Struct Funct. 2018, 2(1):91–109. doi: 10.1007/s00429-017-1478-z.

18. Pellegrino G, Trubert C, Terrien J, Pifferi F, Leroy D, Loyens A, Migaud M, Baroncini M, Maurage CA, Fontaine C, Prévot V, Sharif A. A comparative study of the neural stem cell niche in the adult hypothalamus of human, mouse, rat and gray mouse lemur (Microcebus murinus). J Comp Neurol. 2018, 2(9):1419–1443. doi: 10.1002/cne.24376.

19. Batailler M, Droguerre M, Baroncini M, Fontaine C, Prevot V, Migaud M. DCX-expressing cells in the vicinity of the hypothalamic neurogenic niche: a comparative study between mouse, sheep, and human tissues. J Comp Neurol. 2014, 2(8):1966–85. doi: 10.1002/cne.23514.

20. Knobloch M, Jessberger S. Metabolism and neurogenesis. Curr Opin Neurobiol. 2017, 42:45–52. doi: 10.1016/j.conb.2016.11.006

21. Migaud M, Batailler M, Segura S, Duittoz A, Franceschini I, Pillon D. Emerging new sites for adult neurogenesis in the mammalian brain: a comparative study between the hypothalamus and the classical neurogenic zones. Eur J Neurosci. 2010 Dec;32(12):2042–52. doi: 10.1111/j.1460-9568.2010.07521.x

22. Brus M, Keller M, Lévy F. Temporal features of adult neurogenesis: differences and similarities across mammalian species. Front Neurosci. 2013, 7:135. doi: 10.3389/fnins.2013.00135.

23. Migaud M, Batailler M, Pillon D, Franceschini I, Malpaux B. Seasonal changes in cell proliferation in the adult sheep brain and pars tuberalis. J Biol Rhythms. 2011, 2(6):486–96. doi: 10.1177/0748730411420062.

24. Batailler M, Derouet L, Butruille L, Migaud M. Sensitivity to the photoperiod and potential migratory features of neuroblasts in the adult sheep hypothalamus. Brain Struct Funct. 2016, 2(6):3301–14. doi: 10.1007/s00429-015-1101-0.

25. Chanvallon A, Sagot L, Pottier E, Debus N, François D, Fassier T, Scaramuzzi R.J, Fabre-Nys C. New insights into the influence of breed and time of the year on the response of ewes to the ‘ram effect’. Animal. 2011, 2(10):1594–604. doi: 10.1017/S1751731111000668.

26. Zarazaga LA, Malpaux B, Chemineau P. Amplitude of the plasma melatonin nycthemeral rhythms is not associated with the dates of onset and offset of the seasonal ovulatory activity in the Ile-de-France ewe. Reprod Nutr Dev. 2003, 2(2):167–77. doi: 10.1051/rnd:2003015.

27. Terqui M, Thimonier J. New rapid radioimmunologic method for estimation of plasma progesterone. Application to early diagnosis of gestation in the ewe and goat. C R Acad Hebd Seances Acad Sci D. 1974, 2(13):1109–12.

28. Just N, Adriaensen H, Ella A, Chevillard PM, Batailler M, Dubois JP, Keller M, Migaud M. Blood oxygen level dependent fMRI and perfusion MRI in the sheep brain. Brain Res. 2021, 1760:147390. doi: 10.1016/j.brainres.2021.147390.

29. Gruetter R. Automatic, localized in vivo adjustment of all first-and second-order shim coils. Magn Reson Med. 1993, 2(6):804–11. doi:10.1002/mrm.1910290613.

30. Provencher SW.Estimation of metabolite concentrations from localized in vivo proton NMR spectra. Magn Reson Med. 1993, 2(6):672–9. doi: 10.1002/mrm.1910300604.

31. Tkáč I, Deelchand D, Dreher W, Hetherington H, Kreis R, Kumaragamage C, Považan M, Spielman DM, Strasser B, de Graaf RA. Water and lipid suppression techniques for advanced 1 H MRS and MRSI of the human brain: Experts’ consensus recommendations. NMR Biomed. 2021, 2(5):e4459.doi: 10.1002/nbm.4459.

32. 32Ashburner J. A fast diffeomorphic image registration algorithm. Neuroimage 2007;38:95–113 10.1016/j.neuroimage.2007.07.007

33. Ella A, Delgadillo JA, Chemineau P, Keller M.J Comp Neurol. Computation of a high-resolution MRI 3D stereotaxic atlas of the sheep brain.2017 Feb 15;525(3):676–692. doi: 10.1002/cne.24079.

34. Harris, A.D.; Puts, N.A.; Edden, R.A. Tissue correction for GABA-edited MRS: Considerations of voxel composition, tissue segmentation, and tissue relaxations. J. Magn. Reson. Imaging 2015, 42, 1431–1440.1

35. Near J, Harris AD, Juchem C, Kreis R, Marjańska M, Öz G, Slotboom J, Wilson M, Gasparovic C. Preprocessing, analysis and quantification in single-voxel magnetic resonance spectroscopy: experts’ consensus recommendations.NMR Biomed. 2021 May;34(5):e4257. doi: 10.1002/nbm.4257.

36. 36 Wehr TA. Photoperiodism in humans and other primates: evidence and implications. J Biol Rhythms. 2001, 2(4):348–64. doi: 10.1177/074873001129002060.

37. Malpaux B, Migaud M, Tricoire H, Chemineau P. Biology of mammalian photoperiodism and the critical role of the pineal gland and melatonin. J Biol Rhythms. 2001, 2(4):336–47. doi: 10.1177/074873001129002051.

38. Batailler M, Chesneau D, Derouet L, Butruille L et al. Pineal-dependent increase of hypothalamic neurogenesis contributes to the timing of seasonal reproduction in sheep. Sci Report. 2018, 2(1):6188.doi: 10.1038/s41598-018-24381-4.

39. Dardente H, Wyse CA, Birnie MJ, Dupré SM, Loudon AS, Lincoln GA, Hazlerigg DG. A molecular switch for photoperiod responsiveness in mammals. Curr Biol. 2010, 2(24):2193–8. doi: 10.1016/j.cub.2010.10.048.

40. Nilaweera K, Herwig A, Bolborea M, Campbell G, Mayer CD, Morgan PJ, Ebling FJ, Barrett P. Photoperiodic regulation of glycogen metabolism, glycolysis, and glutamine synthesis in tanycytes of the Siberian hamster suggests novel roles of tanycytes in hypothalamic function. Glia. 2011, 2(11):1695–705. doi: 10.1002/glia.21216.

41. Langlet F. Tanycytes: a gateway to the metabolic hypothalamus. J Neuroendocrinol. 2014, 2(11):753–60. doi: 10.1111/jne.12191.

42. Clark JF, Doepke A, Filosa JA, Wardle RL, Lu A, Meeker TJ, Pyne-Geithman GJ. N-acetylaspartate as a reservoir for glutamate. Med Hypotheses. 2006, 2(3):506–12. doi: 10.1016/j.mehy.2006.02.047.

43. Ryu JR, Hong CJ, Kim JY, Kim EK, Sun W, Yu SW. Control of adult neurogenesis by programmed cell death in the mammalian brain. Mol Brain. 2016, 9:43. doi: 10.1186/s13041-016-0224-4.

44. Beckervordersandforth R. Mitochondrial metabolism-mediated regulation of adult neurogenesis. Brain Plast. 2017, 2(1):73–87. doi: 10.3233/BPL-170044.

45. Chevilllard PM, Batailler M, Piégu B, Estienne A, Blache MC, Dubois JP, Pillon D, Vaudin P, Dupont J, Just N, Migaud M. Seasonal vascular plasticity in the mediobasal hypothalamus of the adult ewe. Histochem Cell Biol. 2022, 2(5):581–593. doi: 10.1007/s00418-022-02079-z.

46. Chevillard PM, Migaud M, Just N. Hypercapnic challenge, BOLD fMRI and immunohistochemistry to examine the in-vivo presence of adult neurogenesis in the sheep hypothalamus. Proceedings of the 2021 annual Meeting of the International Society of Magnetic Resonance in Medicine, p 2945.

47. Zhao X, Ahram A, Berman RF, Muizelaar JP, Lyeth BG. Early loss of astrocytes after experimental traumatic brain injury. Glia, 2003, 44, 140–152. doi: 10.1002/glia.10283

48. Harris JL, Choi IY, Brooks WM. Probing astrocyte metabolism in vivo: proton magnetic resonance spectroscopy in the injured and aging brain. Front Aging Neurosci. 2015, 7:202. doi: 10.3389/fnagi.2015.00202.

49. Moffett JR, Ross B, Arun P, Madhavarao CN, Namboodiri AM. N-Acetylaspartate in the CNS: from neurodiagnostics to neurobiology. Prog Neurobiol. 2007, 2(2):89–131. doi: 10.1016/j.pneurobio.2006.12.003.

50. Ariyannur PS, Arun P, Barry ES, Andrews-Shigaki B, Bosomtwi A, Tang H, Selwyn R, Grunberg NE, Moffett JR, Namboodiri AM. Do reductions in brain N-acetylaspartate levels contribute to the etiology of some neuropsychiatric disorders? J Neurosci Res. 2013, 2(7):934–42. doi: 10.1002/jnr.23234.

51. Lehman MN, Ladha Z, Coolen LM, Hileman SM, Connors JM, Goodman RL. Neuronal plasticity and seasonal reproduction in sheep. Eur J Neurosci. 2010, 2(12):2152–64. doi: 10.1111/j.1460-9568.2010.07530.x.

52. Deelchand DK, Berrington A, Noeske R, Joers JM, Arani A, Gillen J, Schär M, Nielsen JF, Peltier S, Seraji-Bozorgzad N, Landheer K, Juchem C, Soher BJ, Noll DC, Kantarci K, Ratai EM, Mareci TH, Barker PB, Öz G. Across-vendor standardization of semi-LASER for single-voxel MRS at 3T.NMR Biomed. 2021, 2(5):e4218. doi: 10.1002/nbm.4218.

53. Oz G, Alger JR, Barker PB, Bartha R, Bizzi A, Boesch C, Bolan PJ, Brindle KM, Cudalbu C, Dinçer A, Dydak U, Emir UE, Frahm J, González RG, Gruber S, Gruetter R, Gupta RK, Heerschap A, Henning A, Hetherington HP, Howe FA, Hüppi PS, Hurd RE, Kantarci K, Klomp DW, Kreis R, Kruiskamp MJ, Leach MO, Lin AP, Luijten PR, Marjańska M, Maudsley AA, Meyerhoff DJ, Mountford CE, Nelson SJ, Pamir MN, Pan JW, Peet AC, Poptani H, Posse S, Pouwels PJ, Ratai EM, Ross BD, Scheenen TW, Schuster C, Smith IC, Soher BJ, Tkáč I, Vigneron DB, Kauppinen RA; MRS Consensus Group Clinical proton MR spectroscopy in central nervous system disorders.Radiology. 2014, 2(3):658–79. doi: 10.1148/radiol.13130531.

54. Zappaterra MW, Lehtinen MK. The cerebrospinal fluid: regulator of neurogenesis, behavior, and beyond. Cell Mol Life Sci. 2012, 2(17):2863–78. doi: 10.1007/s00018-012-0957-x.

55. Dardente H, Migaud M. Thyroid hormone and hypothalamic stem cells in seasonal functions. Vitam Horm. 2021, 116:91–131. doi: 10.1016/bs.vh.2021.02.005.

56. Eichele G, Bodenschatz E, Ditte Z, Günther AK, Kapoor S, Wang Y, Westendorf C. Cilia-driven flows in the brain third ventricle. Philos Trans R Soc Lond B Biol Sci. 2020? 375 (1792):20190154. doi: 10.1098/rstb.2019.0154.

57. Agnati LF, Fuxe K. Extracellular-vesicle type of volume transmission and tunnelling-nanotube type of wiring transmission add a new dimension to brain neuro-glial networks. Philos Trans R Soc Lond B Biol Sci. 2014, 2(1652):20130505. doi: 10.1098/rstb.2013.0505.

## References

(1) Ella A, Delgadillo JA, Chemineau P, Keller M. Computation of a high-resolution MRI 3D stereotaxic atlas of the sheep brain. Journal of Comparative Neurology. 2017. 525 (3), 676–692

